# Subchondral bone remodeling in knee osteoarthritis is coupled to lipid mobilization from bone marrow adipocytes

**DOI:** 10.64898/2026.10.07.757365

**Authors:** Léa Loisay, Sabrina Bouagga, Diego de Haro, Elke Moradpour, Lorraine Ruffieux, Souad Daamouch, Alex Kanmacher, Thibaut Royon, Alexander Antoniadis, Thomas Hügle, Jeroen Geurts

**Author notes:** Corresponding author: Jeroen Geurts PhD, Service de Rhumatologie, Département de l′appareil locomoteur, Centre Hospitalier Universitaire Vaudois et Université de Lausanne, Avenue Pierre-Decker 4, 1005. These authors contributed equally to this work (Léa Loisay, Sabrina Bouagga). Service of Bone Diseases, Department of Medicine, University Hospital of Geneva, Geneva, Switzerland (Léa Loisay).

## Abstract

**Objective:** Pathological bone formation requires energy derived from metabolic substrates including glucose, amino acids, and fatty acids. As bone marrow adipose tissue (BMAT) is a rich local source of fatty acids and is progressively depleted in osteoarthritic subchondral bone, we investigated whether lipolytic lipid mobilization is associated with osteogenic activity in sclerotic regions and whether this process is pharmacologically modifiable.

**Design:** Human knee osteoarthritis subchondral bone samples (*n* = 23) were analyzed using transcriptomic, histological, and *ex vivo* explant approaches. Associations between lipolytic and bone formation markers were assessed, with particular focus on *PNPLA2*, encoding adipose triglyceride lipase (ATGL), and alkaline phosphatase (ALP). Immunohistochemistry was used to localize lipolytic activation in relation to sclerotic bone regions. *Ex vivo* explants were used to assess adipocyte morphology, secretion of lipolysis products, including free fatty acids (FFA) and glycerol, and bone formation markers including ALP and P1NP. Pharmacological modulation of lipolysis was performed using forskolin to stimulate and metformin to inhibit lipid mobilization.

**Results:** Transcriptomic analysis revealed a strong association between *ALPL* expression and *PNPLA2*, suggesting a link between bone formation and lipolytic activity. Histological analysis confirmed local activation of ATGL in sclerotic subchondral bone regions. In explant cultures, reduced adipocyte size was accompanied by increased release of free fatty acids and glycerol, together with elevated ALP and P1NP levels. FFA secretion strongly correlated with ALP, supporting a relationship between BMAT lipid mobilization and osteogenic activity. *PLIN1*, a marker of lipid droplet integrity, was downregulated in isolated BMAT adipocytes but upregulated in paired stromal-vascular cells, suggesting transfer or utilization of BMAT-derived lipids by neighboring bone-forming cells. Pharmacological modulation confirmed that BMAT lipolysis is dynamically regulated: forskolin enhanced free fatty acid release, whereas metformin reduced it. However, short-term modulation did not significantly alter bone formation indices.

**Conclusion:** These findings identify BMAT lipolysis as a metabolic feature of subchondral bone sclerosis in knee OA. Mobilization of lipids from marrow adipocytes may provide energetic substrates that support osteoblast activity and pathological bone formation. BMAT lipolysis therefore represents a potentially modifiable metabolic pathway within the OA subchondral bone niche.

## Introduction

Bone marrow adipose tissue (BMAT) is increasingly recognized as a dynamic component of the skeletal microenvironment rather than an inert filler of the bone marrow cavity. BMAT responds to nutritional, hormonal, and pathological cues, including caloric restriction and weight loss interventions, and serves as a local reservoir of neutral lipids that can be mobilized during periods of energy demand ^1–5^. This function may be particularly relevant in bone, where osteoblast differentiation, matrix synthesis, and mineralization require energy derived from metabolic substrates including glucose, amino acids, and fatty acids. As BMAT is a rich local source of fatty acids, lipid mobilization from bone marrow adipocytes (BMAd) may contribute to the energetic requirements of bone formation.

Lipolysis is the principal mechanism by which adipocytes release stored lipid-derived energy. Triglycerides contained within lipid droplets are sequentially hydrolyzed by adipose triglyceride lipase (ATGL), encoded by *PNPLA2*, hormone-sensitive lipase (HSL), encoded by *LIPE*, and monoacylglycerol lipase (MGLL), encoded by *MGLL*, resulting in the release of glycerol and free fatty acids ^6^. These fatty acids can be used by neighboring cells as metabolic substrates, including through mitochondrial β-oxidation. In osteoblasts, fatty acid oxidation contributes to postnatal bone acquisition ^7^. Impaired lipid mobilization compromises bone formation, as shown by adipocyte-specific deletion of *Pnpla2*, which reduces parathyroid hormone-induced bone formation and impairs cortical bone repair, and by osteoblast-specific *Pnpla2* deficiency, which reduces cortical and trabecular bone parameters and promotes triglyceride accumulation within bone ^8, 9^. Complementing these findings, conditional adipocyte depletion in lipodystrophic or fat-free mouse models, including BMAds, induces marked bone formation in epiphyseal and diaphyseal bone compartments, further supporting a functional link between marrow adiposity, lipid metabolism, and skeletal anabolism ^10, 11^.

Subchondral bone sclerosis is a hallmark of OA and is characterized by increased bone formation, elevated remodeling activity, and altered tissue mineralization ^12^. These processes are likely to impose high local energy demands. Histologically, sclerotic regions of OA subchondral bone are commonly associated with increased inflammation, vascularity, marrow fibrosis, and a reduction in the number BMAds ^13, 14^. This loss of BMAT has often been interpreted as fibrovascular replacement of marrow fat. However, given the energetic cost of pathological bone formation, an alternative possibility is that BMAds actively mobilize their lipid stores to support local osteogenic activity.

Emerging evidence suggests that BMAT metabolism may contribute to OA-associated bone remodeling. In hand OA, we previously provided the first evidence for BMAT lipolysis in an OA phenotype, showing reduced BMAd size together with increased expression of ATGL, phosphorylated HSL, and MGLL in BMAds and bone-lining cells, as well as correlations between glycerol release and bone formation markers including alkaline phosphatase and P1NP ^15^. However, this work was performed in a distinct OA subtype and mainly revealed general associations between lipolytic and osteogenic markers, without detailed stratification according to subchondral tissue states. In hip OA, transcriptomic profiling further showed that subchondral BMAds are enriched in pathways related to extracellular matrix production and mineralization compared with metaphyseal BMAds, which display a profile more supportive of hematopoiesis ^16^. Together, these observations implicate BMAT metabolism in OA-related bone remodeling, but the role of lipolysis in knee OA subchondral sclerosis remains unexplored.

Therefore, we investigated BMAT lipolysis in paired non-sclerotic and sclerotic regions of human knee OA subchondral bone using transcriptomic, immunohistochemical, and *ex vivo* explant approaches. By assessing changes from non-sclerotic to sclerotic regions within the same patients, this design enabled us to relate local histological measures of BMAd morphology and lipolytic activity to functional explant-based readouts of lipid mobilization and bone formation. We examined whether increased lipolytic activity accompanies osteogenic activation during the transition toward sclerosis and whether this process can be pharmacologically modified. We hypothesized that enhanced BMAT lipolysis supports the energetic demands of osteoblast activity and contributes to subchondral bone sclerosis in OA.

## Methods

### Patient and sample collection

The study protocol was approved by the local ethics committee (CER-VD 2024-0117). Medial and lateral weight-bearing femoral condyles were obtained from consenting patients undergoing elective total knee arthroplasty for primary osteoarthritis (*n* = 23; **Table 1**).

**Table 1.** Demographic characteristics of patients included in the different experimental cohorts.

| **Characteristic** | **Knee condyle**<br>**explants (n = 10)** | **RNA extraction**<br>**(n = 9)** | **Explants with**<br>**treatment (n = 4)** |
| --- | --- | --- | --- |
| Age (years), mean ± SD | 72 ± 9 | 75 ± 11 | 68 ± 12 |
| Female, n (%) | 6 (60%) | 7 (78%) | 2 (50%) |
| BMI (kg/m<sup>2</sup>), mean ± SD | 32.9 ± 7.0 | 27.0 ± 4.0 | 31.0 ± 4.0 |

Specimens were allocated to different experimental workflows. Osteochondral explant cultures combined with histological analyses were performed using samples from 10 patients. In a separate subset of 9 patients, BMAds and stromal vascular fraction (SVF) cells were isolated for RNA extraction and quantitative PCR analysis. Samples from the remaining 4 patients were used for interventional explant experiments.

### Explant culture

Osteochondral explants were prepared from sclerotic and non-sclerotic regions of the weight-bearing femoral condyles using an osteotome and scalpel. Explants were approximately 1 cm in width and 2 mm in thickness. To account for regional variability, explants from each region were cultured in triplicate. Explants were cultured at 37°C and 5% CO_2_ in KRBHA medium, consisting of Krebs-Ringer bicarbonate buffer (Sigma-Aldrich, #K4002) supplemented with HEPES (100 mM; Sigma-Aldrich, #H3537) and fatty acid-free bovine serum albumin (BSA; 0.5% w/v; Sigma-Aldrich, #A7030). For baseline explant experiments, cultures were maintained for 6 days. For interventional experiments, explants were cultured for 7 days in the presence of metformin (100 mM; Sigma-Aldrich, #317240), forskolin (10 mM; Sigma-Aldrich, #E6910), or vehicle control (0.1% v/v DMSO). At the end of the culture period, supernatants were collected and stored at −20°C, while explants were processed for whole-mount staining or fixed in 4% paraformaldehyde for 24 h.

### Whole-mount staining and BMAT area quantification

Osteochondral chips were stained for ALP activity using a solution containing 0.3 mg/mL Naphthol AS-MX phosphate disodium salt (Sigma-Aldrich, #N5000) and 0.3 mg/mL Fast Blue BB salt hemi zinc chloride salt (Sigma-Aldrich, #44670) in 0.1 M Tris-HCl/1 mM MgCl for 5 min. Samples were rinsed with PBS, fixed in 4% paraformaldehyde for 24 h, rinsed again in PBS, and stained with 60% Oil Red O solution (Sigma-Aldrich, #01391) in demineralized water for 5 min. Samples were then briefly washed with PBS.

Whole-mount images of the stained osteochondral chips were acquired using a Leica M205 FA stereomicroscope. Images were analyzed in Fiji to quantify the total tissue area and the Oil Red O-positive area within the bone compartment. The Oil Red O-positive area was used as a measure of BMAT area, and the corresponding fraction was calculated by normalizing this area to the total tissue area.

### Supernatant analysis

Following collection, explant culture supernatants were stored at −20°C until analysis. Adiponectin (R&D Systems, #DY1065-05) and pro-collagen type I N-terminal propeptide (P1NP; R&D Systems, #DY6220-05) concentrations were measured using ELISA kits. Glycerol (Sigma-Aldrich, #MAK266) and free fatty acids (Sigma-Aldrich, #MAK044) were quantified using colorimetric assays. Alkaline phosphatase activity was assessed using p-nitrophenyl phosphate substrate (Sigma-Aldrich, #S0942) in alkaline buffer solution (pH 10.3; Sigma-Aldrich, #A9226) for 2 h at 37°C. Adiponectin, glycerol, and free fatty acid levels were normalized to estimated BMAT mass. Bone-related readouts, including P1NP and alkaline phosphatase activity, were normalized to estimated bone mass.

### Histology and immunohistochemistry

After fixation, osteochondral chips were fully decalcified in 5% formic acid and embedded in paraffin. Serial sections (5 μm) were used for Safranin O, hematoxylin and eosin (H&E), and immunohistochemical staining. Safranin O- and H&E-stained sections were used for Osteoarthritis Bone Score (OABS) scoring, as previously described ^13^. Because no strict histological definition was applied to distinguish sclerotic from non-sclerotic regions, regions were classified based on combined histological and morphological features, including increased bone mass/volume, enhanced ALP staining, reduced BMAT fraction, and frequent association with overlying moderate to severe cartilage degeneration.

For qualitative assessment of protein localization, immunohistochemistry was performed using the Vectastain Elite ABC-HRP kit (Vector Laboratories). Sections were incubated with antibodies against MGLL (1:400; Proteintech, #14986-1-AP), HSL (1:100; Cell Signaling Technology, #4107), phosphorylated HSL (pHSL; 1:100; Cell Signaling Technology, #4139), ATGL (1:100; Cell Signaling Technology, #2138), and Perilipin-1 (PLIN1; 1:200; Cell Signaling Technology, #9349). For HSL, pHSL, ATGL, and PLIN1 staining, antigen retrieval was performed in citrate buffer (pH 6.0) overnight at 60°C. For MGLL staining, antigen retrieval was performed using Tris-EDTA buffer (pH 9.0). Sections were counterstained with Papanicolaou staining. Staining specificity was assessed using secondary antibody-only and isotype IgG controls, as shown in **Supplementary Figure 2**.

Whole-slide images were acquired using a NanoZoomer S60 slide scanner (Hamamatsu). Immunohistochemical stainings were used for qualitative screening and spatial assessment of protein expression. Adipocyte size in sclerotic and non-sclerotic regions was quantified automatically in Fiji using Huang thresholding at 5× magnification within equivalent regions of interest. Across analyzed regions, the number of adipocytes measured per region ranged from 144 to 1,381.

### Stromal vascular fraction and BMAd isolation

BMAds and stromal vascular fraction (SVF) cells were isolated from paired sclerotic and non-sclerotic regions of the femoral condyles. Bone slices were digested in KRBHA medium supplemented with collagenase type II (3 mg/mL; Sigma-Aldrich, #C6885) for 20 min at 37°C under agitation, adapted from the protocol described by Attané *et al.* ^17^.

After digestion, samples were filtered through a 250 μm strainer and centrifuged at 200*g* for 5 min. The floating mature adipocyte fraction was collected, whereas the pelleted SVF fraction was further processed by red blood cell lysis (Sigma-Aldrich, #R7757). Both fractions were collected in TRIzol reagent (Life Technologies, #15596018) and stored at −80°C until downstream RNA extraction.

### RNA extraction and RT-qPCR

BMAd and SVF fractions stored in TRIzol were homogenized using a Precellys Evolution tissue homogenizer (Bertin Technologies) with ceramic beads at 6,000 rpm for two cycles of 5s. For BMAds, the TRIzol phase was collected while avoiding the lipid layer. RNA was extracted and purified using the Direct-zol RNA Miniprep Kit (Zymo Research, #R2052) according to the manufacturer’s instructions. RNA concentration and purity were assessed using a NanoDrop spectrophotometer (Thermo Fisher Scientific).

cDNA was synthesized from 100 ng RNA using SuperScript II reverse transcriptase (Invitrogen, #18064071). RT-qPCR was performed on a QuantStudio 5 Real-Time PCR System (Applied Biosystems) using PowerUp SYBR Green Master Mix (Applied Biosystems, #A25743). Primer sequences are listed in **Supplementary Table 1**. Relative gene expression was calculated using the ΔΔCt method, with the non-sclerotic condition used as the reference.

### Analysis of published bulk transcriptomic datasets

Published bulk transcriptomic datasets from human knee joint tissues were analyzed. Two independent datasets were included: a paired lesional versus non-lesional knee joint dataset comprising samples from 18 donors (EGAS00001004476) ^18^ and a dataset comparing knees with bone marrow lesions (BML; *n* = 14) to non-BML knees (*n* = 6) (E-MTAB-5564) ^19^.

A curated set of genes was selected based on established roles in BMAT lipolysis, fatty acid metabolism, and bone formation ^20^. To account for differences in adipocyte content between samples, expression values were re-normalized using an adipocyte-content score, calculated as the average expression of five adipocyte-enriched marker genes (*PLIN1*, *GPD1*, *ADIPOQ*, *CIDEC*, *PLAAT3*), as previously described ^21^. Pairwise correlations between the normalized expression values of selected genes were calculated within each dataset. The resulting correlation coefficients were hierarchically clustered and visualized as heatmaps to compare gene–gene association patterns across datasets.

### Statistical methods

No formal a priori sample size calculation was performed because this exploratory study relied on the availability of human osteochondral specimens obtained from patients undergoing joint replacement surgery. Sample size was determined by tissue availability and prior experience with paired human osteochondral explant analyses ^22^. The paired study design, in which sclerotic and non-sclerotic regions were obtained from the same donor, was used to reduce inter-individual variability, and paired statistical analyses were performed wherever possible. Data distribution was assessed using the Anderson–Darling, Shapiro– Wilk, Kolmogorov–Smirnov, and D’Agostino–Pearson normality tests. For paired comparisons, normally distributed data were analyzed using paired t-tests, whereas non-normally distributed data were analyzed using Wilcoxon matched-pairs signed-rank tests. For analyses involving multiple explants obtained from the same donor, mixed-effects models were used with donor included as a random effect to account for within-donor clustering and the non-independence of observations. Associations between continuous variables were assessed using Pearson correlation analysis after confirming normality. Individual data points are shown in the figures, with confidence intervals reported where applicable. Exact p values are reported in the figure legends. Asterisks in figure panels are used only as visual indicators of statistically significant comparisons, with p < 0.05 considered statistically significant. All statistical analyses were performed using GraphPad Prism version 10.6.0 (GraphPad Software, Boston, MA, USA).

## Results

### BMAT lipid metabolism associates with bone formation signatures in knee OA subchondral bone

We previously observed increased expression of lipolysis-associated proteins in regions of pathological bone formation in hand OA ^15^. To determine whether a similar association may occur in knee OA subchondral bone, we analyzed two published bulk transcriptomic datasets from human knee joint tissues, including samples stratified by bone marrow lesion status ^19^ and paired preserved/lesional regions ^18^.

Pairwise correlations were calculated between selected genes involved in lipolysis, lipid droplet biology, fatty acid metabolism, and bone formation. Hierarchical clustering of the resulting correlation coefficients revealed coordinated relationships between lipid metabolism and bone formation-associated genes across datasets (**Fig. 1a**). *ALPL*, *COL1A1*, and *COL1A2* correlated with fatty acid metabolism genes, including *SCD1*, *FADS1*, and *FADS2*, and with lipolysis-associated genes such as *PNPLA2* and *ABHD5*. In contrast, the lipolysis inhibitor *G0S2* showed an opposing association pattern. Accordingly, *PNPLA2* expression was inversely correlated with *G0S2* and positively correlated with *ALPL* (**Fig. 1b**). These data suggest that knee OA subchondral bone exhibits coordinated transcriptional relationships between BMAT lipid mobilization and bone formation-associated remodeling.

**Figure 1.**
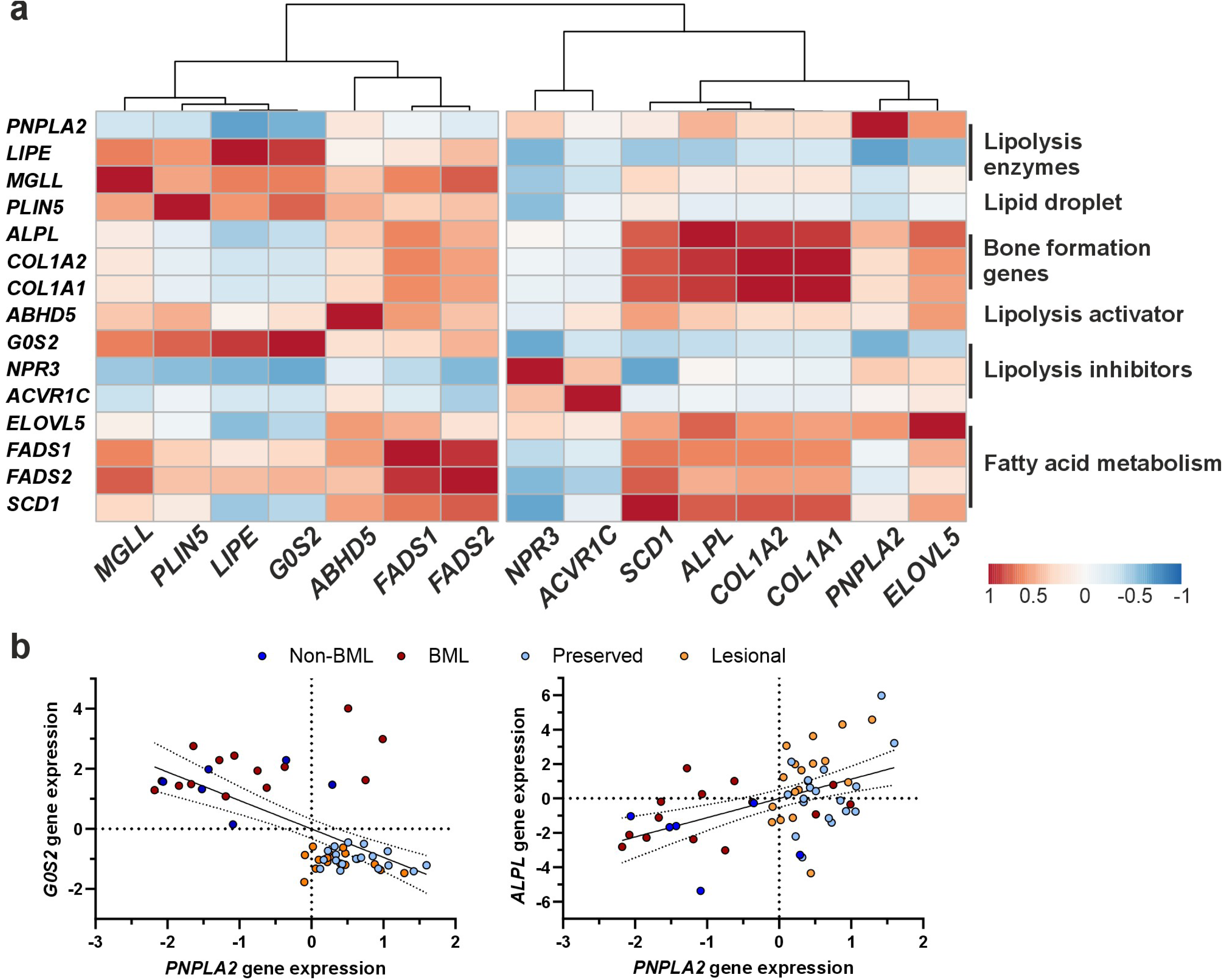
Lipolysis and fatty acid metabolism are associated with osteogenic gene expression in human OA subchondral bone. (**a**) Hierarchically clustered heatmap of Pearson correlation coefficients between genes involved in lipolysis, lipid-droplet regulation, bone formation, regulation of lipolysis, and fatty acid metabolism in two published transcriptomic datasets of human knee osteoarthritic subchondral bone. Rows and columns show the same genes, grouped according to their proposed biological functions. Red indicates positive and blue indicates negative correlations, with color intensity representing correlation strength. (**b**) Associations between *PNPLA2* expression and expression of the endogenous ATGL inhibitor *G0S2* (left) or the osteogenic marker *ALPL* (right). Samples are colored according to bone marrow lesion status in one dataset and preserved or lesional tissue classification in the second dataset. Solid lines represent linear regression fits, and dotted lines indicate 95% confidence intervals. Vertical and horizontal dotted lines indicate zero-centered gene-expression values. ATGL, adipose triglyceride lipase; BML, bone marrow lesion; ALPL, alkaline phosphatase; G0S2, G0/G1 switch gene 2; LIPE, hormone-sensitive lipase; MGLL, monoglyceride lipase; PNPLA2, patatin-like phospholipase domain-containing protein 2.

### Sclerotic subchondral bone in knee OA shows reduced BMAT area and smaller BMAds

We next characterized paired osteochondral explants from human OA femoral condyles to define the tissue context in which BMAT remodeling occurs. Non-sclerotic and sclerotic regions were analyzed in parallel from the same donor. Regional classification was based on subchondral bone remodeling, supported by OABS scoring shown together with the corresponding histological images in **Supplementary Table 2 and Fig. 1**.

Whole-mount staining illustrated clear regional differences between paired explants. Non-sclerotic regions showed limited alkaline phosphatase activity and abundant Oil Red O-positive marrow adipose area, whereas sclerotic regions showed stronger alkaline phosphatase staining, increased bone area, and visibly reduced Oil Red O-positive BMAT (**Fig. 2a**). Quantitative image analysis showed that BMAT fraction decreased as bone fraction increased across the analyzed explants (**Fig. 2b**), consistent with expansion of the bone compartment being accompanied by a proportional reduction of the marrow adipose compartment.

**Figure 2.**
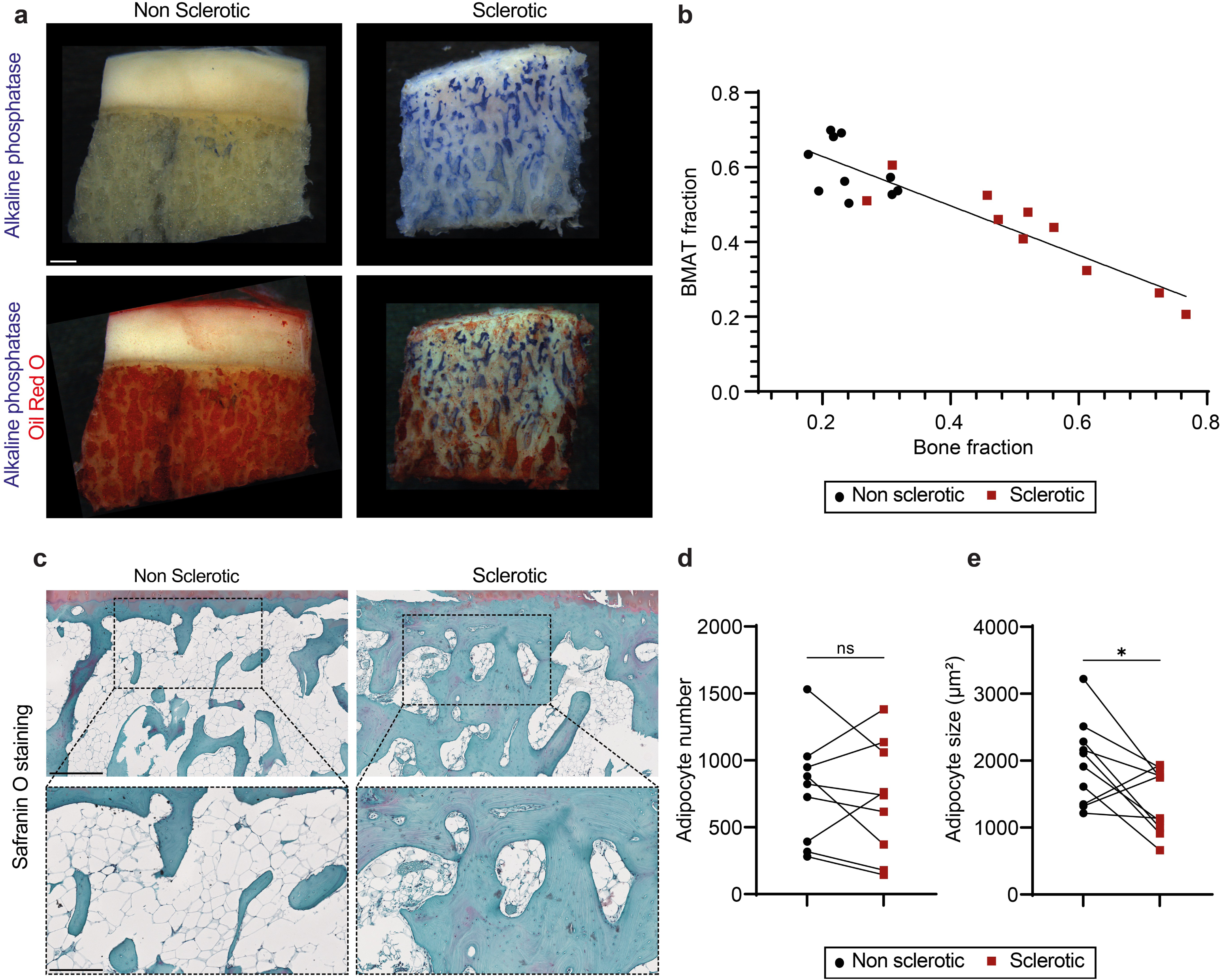
Bone marrow adipose tissue is reduced in sclerotic OA subchondral bone. (**a**) Representative whole-mount images of paired non-sclerotic and sclerotic osteochondral explants stained for alkaline phosphatase activity, either alone or in combination with Oil Red O to visualize BMAT. Scale bar, 1 mm. (**b**) Association between BMAT fraction and bone fraction across non-sclerotic and sclerotic explants. Symbols represent individual explants, and the solid line represents the linear regression fit. (**c**) Representative Safranin O-stained sections of paired non-sclerotic and sclerotic osteochondral regions. Dashed boxes indicate the regions shown at higher magnification. Scale bars, 500 µm in the upper panels and 250 µm in the lower panels. (**d,e**) Paired quantification of BMAd number (**d**) and mean BMAd size (**e**) in non-sclerotic and sclerotic regions. Each connected pair represents samples from the same donor. Black circles indicate non-sclerotic regions and red squares indicate sclerotic regions. * *P* < 0.05; ns, not significant.

Safranin O-stained sections further supported the regional characterization of the explants, with sclerotic regions showing increased subchondral bone and reduced marrow adipose space compared with paired non-sclerotic regions (**Fig. 2c**). To determine whether the reduced BMAT area reflected adipocyte depletion or altered adipocyte morphology, BMAds were quantified in matched regions of interest. BMAd number did not differ consistently between non-sclerotic and sclerotic regions (**Fig. 2d**). In contrast, BMAd size was reduced in sclerotic regions (**Fig. 2e**). BMAd size in non-sclerotic regions was not associated with patient BMI (Pearson *r* = 0.28, 95% CI −0.42 to 0.77), suggesting that baseline BMAd morphology was not primarily explained by systemic adiposity.

### Sclerotic subchondral bone shows coordinated changes in lipolysis products and bone formation markers

Culture supernatants from paired non-sclerotic and sclerotic explants were analyzed to determine whether local tissue remodeling was accompanied by altered secretory activity. Sclerotic explants released higher levels of ALP activity and P1NP than paired non-sclerotic explants, consistent with increased bone formation-associated activity in remodeled regions (**Fig. 3a,b**). Sclerotic explants also released higher levels of glycerol and free fatty acids, both products of lipolysis, as well as increased adiponectin when normalized to BMAT mass (**Fig. 3c–e**). Thus, despite reduced BMAT area and smaller BMAds, sclerotic regions displayed increased adipose-associated secretory activity.

**Figure 3.**
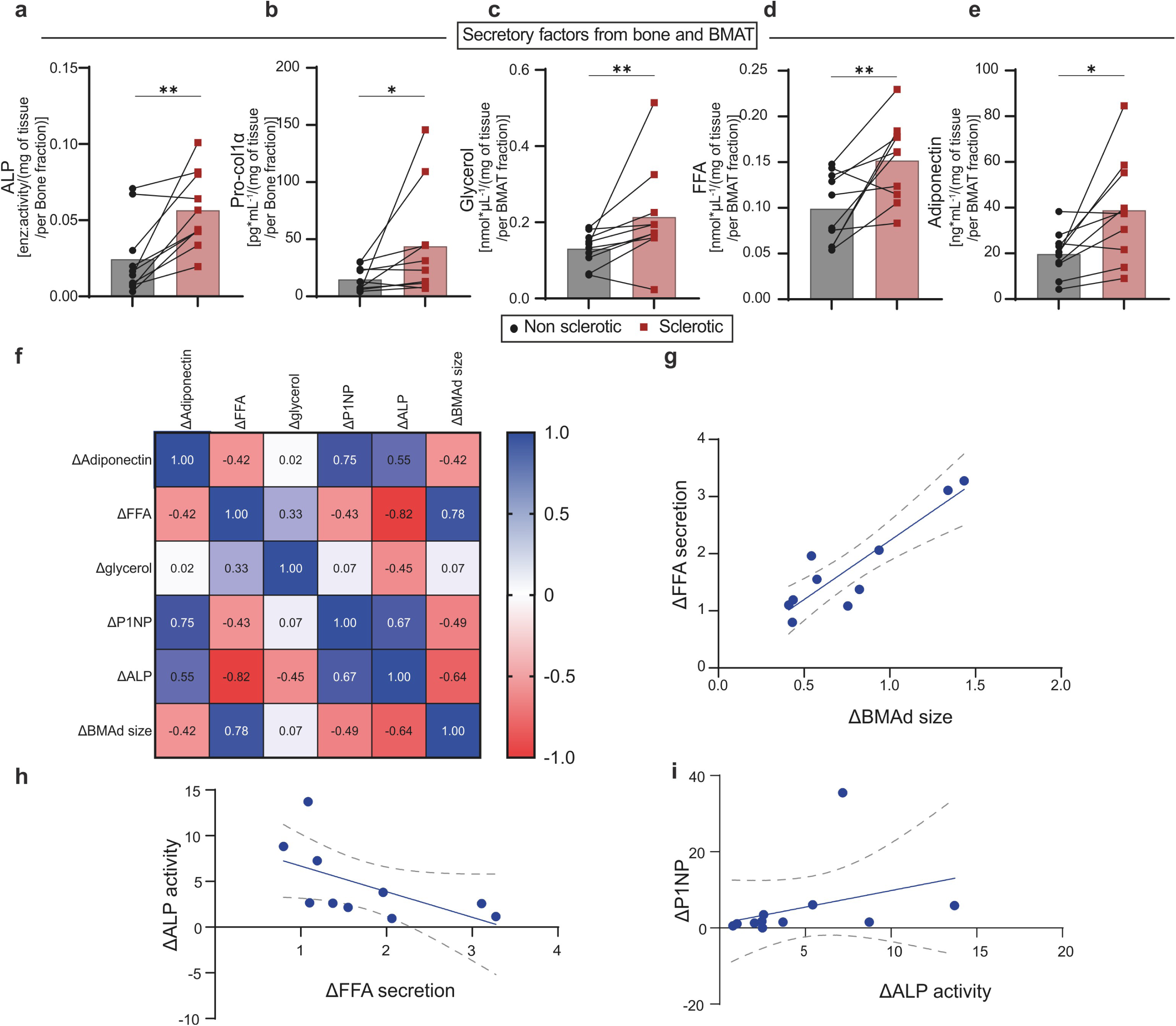
Increased lipid mobilization is associated with bone formation in sclerotic OA subchondral bone. Secretory activity of paired non-sclerotic and sclerotic osteochondral explants and associations between regional changes in BMAT and bone formation. (**a,b**) Bone formation-associated readouts in explant culture supernatants: alkaline phosphatase activity normalized to estimated bone mass (**a**) and procollagen type I N-terminal propeptide normalized to estimated bone mass (**b**). (c–e) BMAT-associated secretory readouts: glycerol release (**c**), free fatty acid secretion (**d**), and adiponectin secretion (**e**), normalized to estimated BMAT mass. Bars represent group means, symbols represent individual explants, and connecting lines identify paired non-sclerotic and sclerotic samples from the same donor. Black circles indicate non-sclerotic regions and red squares indicate sclerotic regions. (**f**) Pearson correlation matrix showing associations between paired regional differences, calculated as sclerotic minus non-sclerotic values, in adiponectin secretion, FFA secretion, glycerol release, P1NP secretion, ALP activity, and mean BMAd size. Numbers indicate Pearson correlation coefficients. Blue indicates positive correlations and red indicates negative correlations. (**g-i**) Associations between paired regional differences in BMAd size and FFA secretion (**g**), FFA secretion and ALP activity (**h**), and ALP activity and P1NP secretion (**i**). Associations were assessed using Pearson correlation. Solid lines represent linear regression fits, and dashed lines indicate 95% confidence intervals. Δ indicates the difference between paired sclerotic and non-sclerotic regions. *P < 0.05; **P < 0.01.

To assess whether these changes were coordinated within donors, fold changes from non-sclerotic to sclerotic regions were calculated for each explant pair. Correlation analysis of these fold changes revealed associations between BMAd size, lipolytic products, and bone formation-associated readouts (**Fig. 3f**). Selected correlations showed that the fold-change increase in FFA release was associated with the change in BMAd size and inversely associated with the fold-change increase in ALP activity, whereas ALP activity and P1NP secretion were positively associated (**Fig. 3g–i**). Although FFA release was increased overall in sclerotic explants, samples with the strongest increase in ALP activity showed a lower relative accumulation of FFA in the supernatant. This suggests that measured FFA levels reflect not only lipolytic release, but also local fatty acid uptake, retention, or utilization by highly remodeling tissue.

### Lipolysis-associated proteins show spatially restricted patterns in sclerotic subchondral bone

Because sclerotic explants released increased levels of glycerol and free fatty acids, we next assessed whether canonical lipolysis-associated proteins were detectable in BMAds and surrounding remodeling areas. Immunohistochemistry was used as a qualitative approach to evaluate the spatial distribution of Perilipin-1, ATGL, HSL, phosphorylated HSL, and MGLL in non-sclerotic and sclerotic subchondral bone. Perilipin-1 staining delineated lipid droplets in BMAds in both non-sclerotic and sclerotic regions, with smaller Perilipin-1-positive BMAds apparent in sclerotic tissue, consistent with the reduced BMAd size observed histologically (**Fig. 4a**). HSL and MGLL were detectable in BMAds in non-sclerotic regions, whereas ATGL staining was weak or limited (**Fig. 4b,c,e**). In sclerotic regions, ATGL, phosphorylated HSL, and MGLL staining were more apparent in BMAds and in stromal or fibrotic areas adjacent to remodeled bone surfaces (**Fig. 4b,d,e**). Comparable staining patterns across additional sclerotic regions are shown in **Supplementary Fig. 3 & 4**. These observations suggest that lipolysis-associated proteins are not uniformly distributed throughout the tissue but are instead enriched in specific cellular compartments near areas of active remodeling.

**Figure 4.**
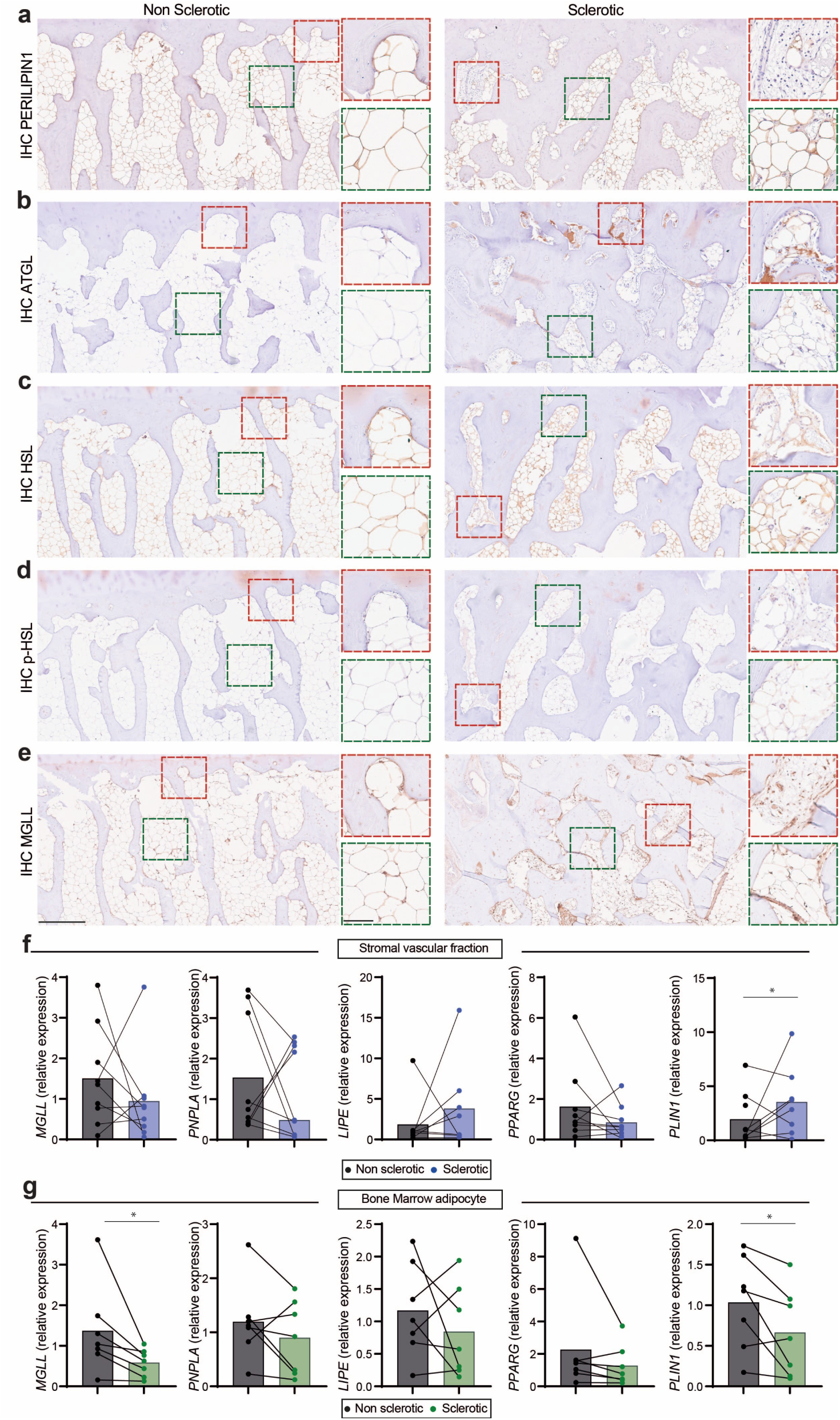
Lipolytic enzymes are differentially localized and expressed in sclerotic osteoarthritic subchondral bone. Representative immunohistochemical staining of paired non-sclerotic and sclerotic subchondral bone for **(a)** Perlipin1, **(b)** adipose triglyceride lipase (ATGL), **(c)** hormone-sensitive lipase (HSL), **(d)** phosphorylated HSL (p-HSL), and **(e)** monoglyceride lipase (MGL). Red- and green-dashed boxes identify regions shown at higher magnification. **(f, g)** Relative expression of *MGLL*, *PNPLA2*, *LIPE*, *PPARG*, and *PLIN1* in the stromal vascular fraction **(f)** and isolated bone marrow adipocytes **(g)** from paired non-sclerotic and sclerotic regions. Bars represent group means, symbols represent individual samples, and connecting lines identify paired samples from the same donor. Black circles indicate non-sclerotic regions, blue circles indicate the stromal vascular fraction from sclerotic regions, and green circles indicate bone marrow adipocytes from sclerotic regions. Paired comparisons were performed between non-sclerotic and sclerotic regions. \**P* < 0.05.

To determine whether these spatial patterns were reflected at the transcript level, lipolysis-related genes were analyzed separately in isolated stromal vascular fraction and BMAd fractions. In the stromal vascular fraction, expression patterns were variable across donors, but *PLIN1* was increased in sclerotic regions (**Fig. 4f**). In contrast, *PLIN1* was reduced in isolated BMAds from sclerotic regions, consistent with smaller mature adipocytes and reduced lipid droplet content (**Fig. 4g**). *MGLL* expression was also reduced in the BMAd fraction, whereas other lipolysis-associated genes showed no consistent directional change.

Together, these findings suggest compartment-specific remodeling of lipid droplet and lipolysis-associated pathways in sclerotic subchondral bone. The qualitative protein localization highlights spatial patterns near remodeling areas that are not fully captured by bulk gene expression in isolated fractions.

### Lipolytic output can be modulated in osteochondral explants

Finally, we used pharmacological stimulation and inhibition of lipolysis to assess whether FFA release in the osteochondral explant model reflected a dynamic lipolytic process. Paired non-sclerotic and sclerotic explants were cultured for 7 days with vehicle control, forskolin, or metformin. Whole-mount ALP staining after culture showed that regional tissue characteristics were maintained during treatment, with sclerotic explants retaining stronger ALP-positive staining than non-sclerotic explants across conditions (**Fig. 5a**). In non-sclerotic explants, forskolin increased FFA release, whereas metformin reduced FFA release compared with vehicle control (**Fig. 5b**). These changes were not accompanied by detectable changes in ALP activity over the same culture period (**Fig. 5c**). In sclerotic explants, FFA release remained elevated and variable across treatment conditions, without a consistent treatment-associated shift in FFA release or ALP activity (**Fig. 5d,e**).

**Figure 5.**
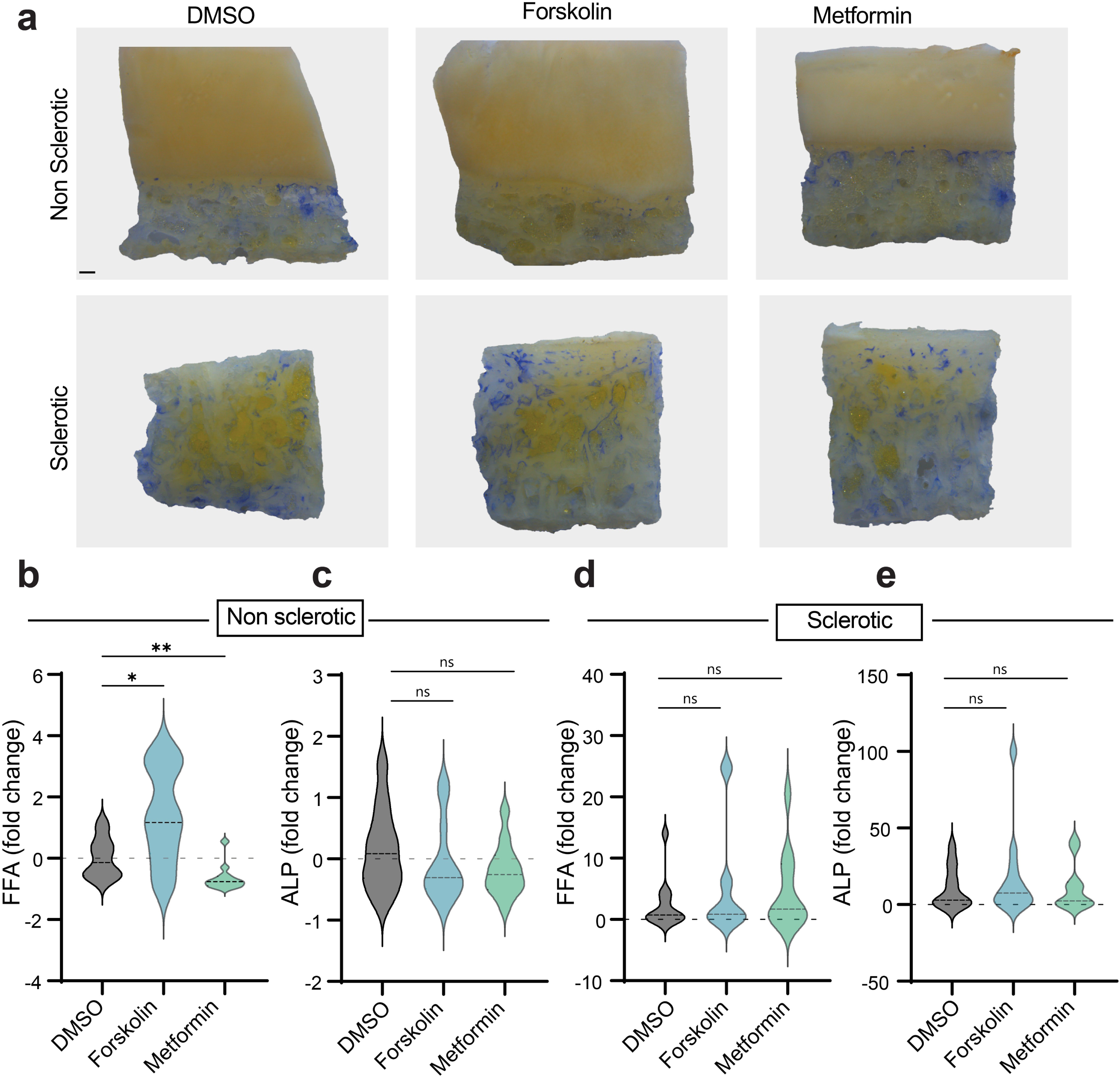
Pharmacological modulation of lipolysis differentially affects non-sclerotic and sclerotic OA subchondral bone. (**a**) Representative whole-mount images of alkaline phosphatase-stained non-sclerotic and sclerotic osteochondral explants treated with vehicle control (DMSO), forskolin, or metformin. (**b,c**) Treatment-to-control fold changes in free fatty acid secretion (**b**) and alkaline phosphatase activity (**c**) in non-sclerotic explants. Forskolin increased FFA secretion, whereas metformin reduced FFA secretion, without significantly affecting ALP activity. (**d,e**) Treatment-to-control fold changes in FFA secretion (**d**) and ALP activity (**e**) in sclerotic explants. Values were normalized to the corresponding DMSO-treated control explants from the same donor. Violin plots show the distribution of individual explant responses, and dashed horizontal lines indicate the median. Statistical comparisons were performed against the corresponding DMSO control. *P < 0.05; **P < 0.01; ns, not significant.

These findings indicate that FFA release from osteochondral explants reflects a modifiable lipolytic process. However, under the present short-term culture conditions, changes in FFA release were not accompanied by detectable changes in ALP activity.

## Discussion

Subchondral sclerosis is characterized by accelerated bone remodeling and formation, yet how the energetic demands of this process are met remains poorly understood. Our findings identify BMAT lipolysis as a potential component of this response. In sclerotic regions, BMAds were smaller but not fewer, while glycerol and FFA release, lipolytic activity, and expression of genes involved in fatty acid metabolism were increased. The spatial localization of lipolytic proteins near active bone-forming surfaces, together with the association between FFA release and ALP and P1NP, suggests that lipid mobilization is coupled to pathological bone formation. These observations support a model in which BMAT is not progressively lost during OA but is metabolically reprogrammed to release stored lipid within remodeling niches.

Evidence from experimental models indicates that BMAd lipolysis is largely dispensable during skeletal homeostasis but becomes important when bone loss, repair, or sharply increased bone formation creates additional energetic demand. BMAd-specific deletion of *Pnpla2*, encoding ATGL, has limited skeletal consequences under normal feeding conditions but compromises bone maintenance during caloric restriction and cold exposure and impairs bone regeneration after injury ^9^. Similarly, the anabolic response to intermittent PTH requires ATGL-dependent fatty acid release from adipocytes and subsequent fatty acid oxidation by osteoblasts ^23^. Sclerotic OA bone, with its persistently elevated turnover and remodeling, may represent an analogous state in which local lipid stores become functionally important. Consistent with this context dependency, forskolin increased FFA release from non-sclerotic explants without affecting bone formation indices. Thus, lipolysis can be induced in non-sclerotic bone, but lipid release alone may be insufficient to stimulate osteogenesis. Additional features of the sclerotic environment appear necessary to couple lipid mobilization to bone formation.

The metabolic contribution of BMAT may operate alongside its paracrine regulation of osteogenesis. Ablation of adiponectin-expressing cells causes extraordinarily rapid BMPR-dependent bone formation through depletion of the BMP inhibitors GREM1 and CHRDL1, with bone mass increasing approximately tenfold within 10 to 14 days ^11^. This study establishes that marrow adipose-lineage cells can restrain osteogenesis but does not address the fate of their lipid stores or how the energetic demands of such rapid bone formation are supported. Moreover, osteoblasts themselves contain lipid droplets and require ATGL-dependent intracellular lipolysis, particularly during later stages of differentiation and mineralization ^8^. Lipolysis in sclerotic bone may therefore involve both the release of fatty acids by BMAds and the mobilization of intracellular lipid within osteoblast-lineage cells. Our data cannot yet distinguish these processes or demonstrate direct transfer of BMAd-derived fatty acids to osteoblasts, but they indicate that lipid mobilization occurs at the interface between BMAT and active bone formation.

An important unresolved question is how this lipolytic response is activated in the loaded OA joint. IL-11 provides a plausible link because it is produced by osteoblasts and osteocytes in response to mechanical loading and regulates both osteogenesis and adipose metabolism ^24^. Notably, IL11 was also among the prominent genes identified in lesional OA subchondral bone by Tuerlings and coworkers ^18^. A second candidate is RCN2, which is secreted by marrow macrophages in response to mechanical stimulation in mice and activates cAMP-PKA-dependent BMAT lipolysis to support osteogenesis ^25^. Although this mechanism has not been validated in human tissue, macrophages have previously been observed between BMAds and in crown-like arrangements in OA bone, placing them in an appropriate location to regulate lipid mobilization ^26^. Our reanalyzed transcriptomic data further suggest that control of ATGL activity may be important. Reduced G0S2, an endogenous ATGL inhibitor, combined with increased ABHD5, an ATGL activator, could promote lipolysis without requiring increased ATGL expression. However, the signals regulating this balance in OA, and whether they arise from loading, inflammation, macrophages, or osteoblast-lineage cells, remain unknown.

These findings also raise questions about the effects of metabolic therapies on the OA joint. Forskolin confirms that cAMP-sensitive lipolysis can be activated in human subchondral tissue, but forskolin and metformin act on several cell types and cannot establish that their effects on bone are mediated specifically through BMAT. More selective approaches will be needed to determine whether pharmacological modulation of lipolysis alters FFA supply to osteoblasts or acts directly on osteoblast metabolism. This issue is clinically relevant because semaglutide substantially reduced pain in people with obesity and knee OA in the STEP 9 trial, although the relative contributions of weight loss, reduced joint loading, and direct tissue effects remain unresolved ^27^. In a separate population with increased fracture risk, semaglutide increased CTX without increasing P1NP and was associated with lower hip and spine BMD relative to placebo ^28^. Systemic turnover markers and measurements from the axial skeleton, however, may not predict the response of highly remodeled sclerotic subchondral bone. Whether metabolic drugs modify BMAT lipolysis or subchondral remodeling independently of their effects on body weight and joint loading is therefore an important question.

Several limitations temper our interpretation. The cross-sectional analysis of end-stage OA tissue cannot establish whether lipolysis drives sclerosis or develops in response to increased bone formation. Reduced BMAd size and increased FFA and glycerol release support lipid mobilization, but direct transfer and oxidation of BMAd-derived fatty acids by osteoblasts were not demonstrated. Lipolytic proteins in bone-lining cells also indicate that osteoblast-intrinsic lipolysis may contribute independently of BMAT. In addition, our pharmacological interventions were not cell selective, and the proposed roles of IL-11, RCN2, macrophages, and the G0S2-ABHD5-ATGL axis require direct validation in human OA tissue. Nevertheless, our findings support a context-dependent metabolic switch in which abnormal loading and the sclerotic microenvironment couple local lipolysis to pathological bone formation. The therapeutic objective may therefore not be to suppress lipolysis broadly, but to normalize maladaptive lipid mobilization without compromising the fatty acid utilization required for physiological remodeling and repair.

## Acknowledgements

The authors have no additional contributors to acknowledge.

## Contributions

Léa Loisay, Sabrina Bouagga, Thomas Hügle, and Jeroen Geurts contributed to the conception and design of the study. Léa Loisay, Sabrina Bouagga, Alexander Antoniadis, Thomas Hügle, and Jeroen Geurts contributed to the experimental design. Léa Loisay, Sabrina Bouagga, Diego de Haro, Elke Moradpour, Lorraine Ruffieux, Thibaut Royon, Alexander Antoniadis, Souad Daamouch, and Alex Kanmacher contributed to the acquisition and assembly of data. Léa Loisay, Sabrina Bouagga, Diego de Haro, Elke Moradpour, Alexander Antoniadis, Thomas Hügle, and Jeroen Geurts contributed to the analysis and interpretation of data. Léa Loisay, Sabrina Bouagga, and Jeroen Geurts drafted the manuscript. All authors critically revised the manuscript for important intellectual content, approved the final version for publication, and agree to be accountable for all aspects of the work.

Jeroen Geurts takes responsibility for the integrity of the work as a whole, from inception to finished article.

## Declaration of Generative AI and AI-assisted technologies in the writing process

During the preparation of this work, the authors used Microsoft 365 Copilot to improve the readability and language of the manuscript. After using this tool, the authors reviewed and edited the content as needed and take full responsibility for the content of the publication.

## Author contributions

Conceptualization: Loisay, Bouagga, Hügle, Geurts

Experimental design: Loisay, Bouagga, Antoniadis, Hügle, Geurts

Data acquisition: Loisay, Bouagga, de Haro, Moradpour, Ruffieux, Royon,, Antoniadis

Data analysis and interpretation: Loisay, Bouagga, de Haro, Moradpour, Daamouch, Kanmacher, Antoniadis, Hügle, Geurts

Writing: Loisay, Bouagga, Geurts

