## Supplementary Tables and Figures for "Subchondral bone remodeling in knee osteoarthritis is coupled to lipid mobilization from bone marrow adipocytes"

Running title: Bone marrow adipocyte lipolysis in knee OA

Léa Loisay, Sabrina Bouagga, Diego de Haro, Elke Moradpour, Lorraine Ruffieux, Souad Daamouch, Alex Kanmacher, Thibaut Royon, Alexander Antoniadis, Thomas Hügle, Jeroen Geurts

**Data Supplement**

**Supplementary Table 1. Primer sequences used for RT-qPCR**

| Gene name | Primer sequence (5`→3`) |
| --- | --- |
| *GAPDH* | FW – TCCTGCACCACCAACTGCTTAG  RV – TGGACTGTGGTCATGAGTCCTTC |
| *MGLL* | FW – GGCATGGTACTCATTTCGCCTC  RV – GTTTGGCAGCACAAGGTTGAGC |
| *PNPLA2* | FW – CCCACTTCAACTCCAAGGACG  RV – GCAGGTTGTCTGAAATGCCACC |
| *LIPE* | FW – AGCCTTCTGGAACATCACCGAG  RV –TCGGCAGTCAGTGGCATCTCAA |
| *PLIN1* | FW – GACAACGTGGTGGACACAGT  RV – CTGGTGGGTTGTCGATGTC |

**Supplementary Table 2. Individual Osteoarthritis Bone Score (OABS) components and total scores in matched non-sclerotic and sclerotic osteochondral regions**

A. Non-sclerotic regions

| OABS component | 107 | 109 | 110 | 111 | 114 | 116 | 119 | 120 | 121 | 122 |
| --- | --- | --- | --- | --- | --- | --- | --- | --- | --- | --- |
| Cysts | 0 | 0 | 0 | 0 | 0 | 0 | 0 | 0 | 0 | 0 |
| Fibrosis | 1 | 1 | 1 | 1 | 1 | 1 | 1 | 1 | 1 | 1 |
| Blood vessels | 1 | 1 | 1 | 1 | 1 | 1 | 1 | 1 | 0 | 0 |
| Cartilage islands | 0 | 0 | 0 | 0 | 1 | 0 | 0 | 1 | 0 | 0 |
| Thickened trabeculae | 0 | 0 | 1 | 0 | 0 | 1 | 0 | 0 | 0 | 0 |
| Tidemark integrity | 0 | 0 | 1 | 1 | 1 | 1 | 1 | 1 | 1 | 1 |
| Inflammation | 0 | 1 | 1 | 1 | 0 | 1 | 1 | 1 | 1 | 1 |
| Total OABS score | **2** | **3** | **5** | **4** | **4** | **5** | **4** | **5** | **3** | **3** |

B. Matched sclerotic regions

| OABS component | 107 | 109 | 110 | 111 | 114 | 116 | 119 | 120 | 121 | 122 |
| --- | --- | --- | --- | --- | --- | --- | --- | --- | --- | --- |
| Cysts | 0 | 0 | 0 | 1 | 0 | 1 | 0 | 0 | 1 | 0 |
| Fibrosis | 1 | 1 | 1 | 1 | 1 | 1 | 1 | 1 | 1 | 1 |
| Blood vessels | 1 | 1 | 1 | 1 | 1 | 1 | 1 | 1 | 1 | 1 |
| Cartilage islands | 0 | 0 | 0 | 0 | 0 | 1 | 0 | 0 | 1 | 0 |
| Thickened trabeculae | 1 | 1 | 1 | 1 | 0 | 1 | 1 | 1 | 1 | 1 |
| Tidemark integrity | 1 | 1 | 1 | 1 | 0 | 1 | 1 | 1 | 1 | 1 |
| Inflammation | 1 | 0 | 1 | 1 | 1 | 1 | 1 | 1 | 1 | 1 |
| Total OABS score | **5** | **4** | **5** | **6** | **3** | **7** | **5** | **5** | **7** | **5** |

Histological features were scored as absent (0) or present (1). Total OABS scores were calculated as the sum of the seven component scores, ranging from 0 to 7. Non-sclerotic and sclerotic regions sharing the same anonymized donor identifier represent matched osteochondral regions obtained from the same donor.

**Supplementary Figure 1. Safranin-*O* staining of osteochondral samples included in the baseline secretion experiments**

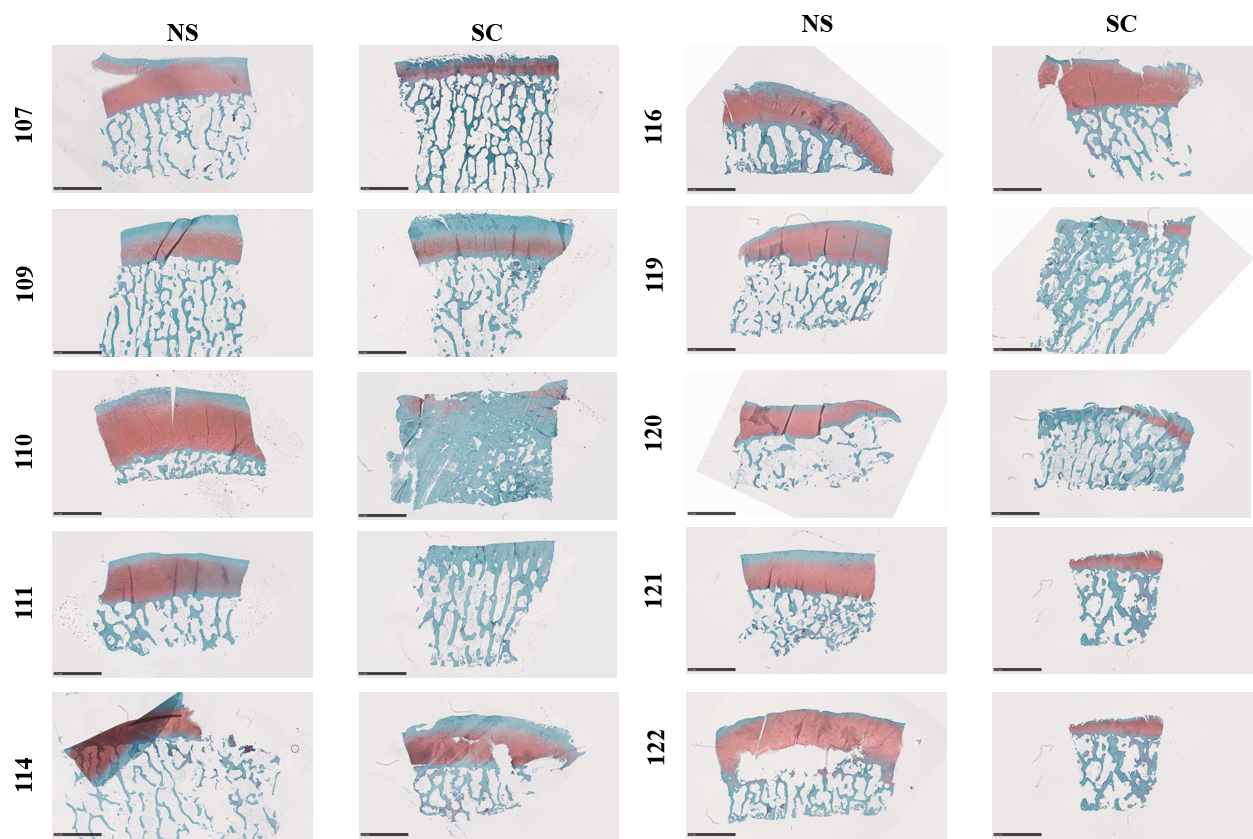

Safranin O-stained sections of matched non-sclerotic and sclerotic osteochondral regions from all donors included in the baseline secretion analyses. Sample identifiers correspond to the anonymized donor identifiers used throughout the supplementary material. Scale bars: 2.5 mm

**Supplementary Figure 2. Antibody controls for immunohistochemical detection of lipolysis-associated proteins in human subchondral bone**

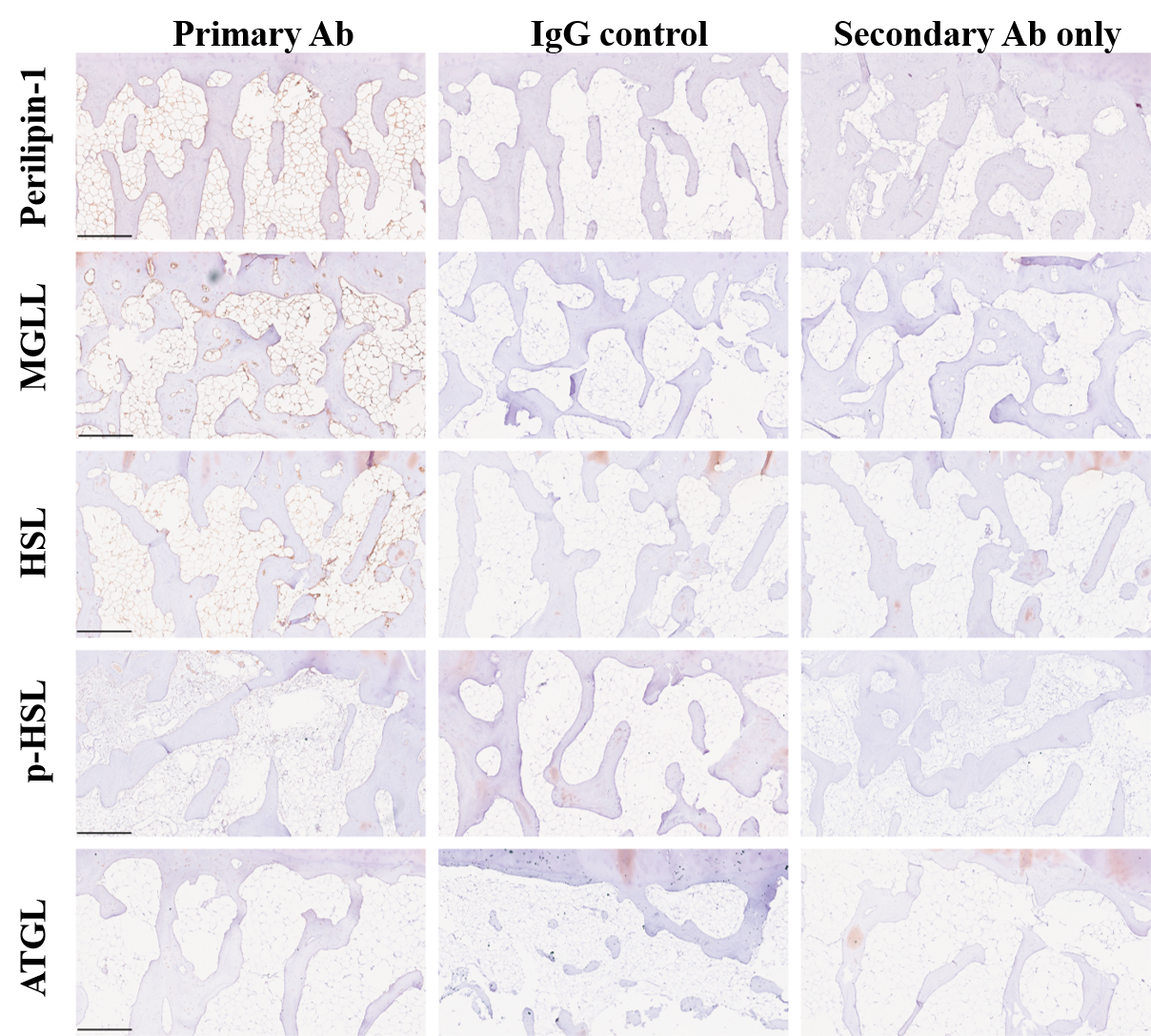

Representative immunohistochemical staining for Perilipin-1, MGLL, HSL, phosphorylated HSL (p-HSL), and ATGL in human osteochondral tissue. Sections incubated with the respective primary antibody are shown alongside sections incubated with the corresponding IgG control antibody or processed with secondary antibody only. Brown staining indicates immunoreactivity, with hematoxylin counterstaining. Scale bars: 500 μm.

**Supplementary Figure 3. Immunohistochemical staining for ATGL and MGLL in sclerotic subchondral bone**

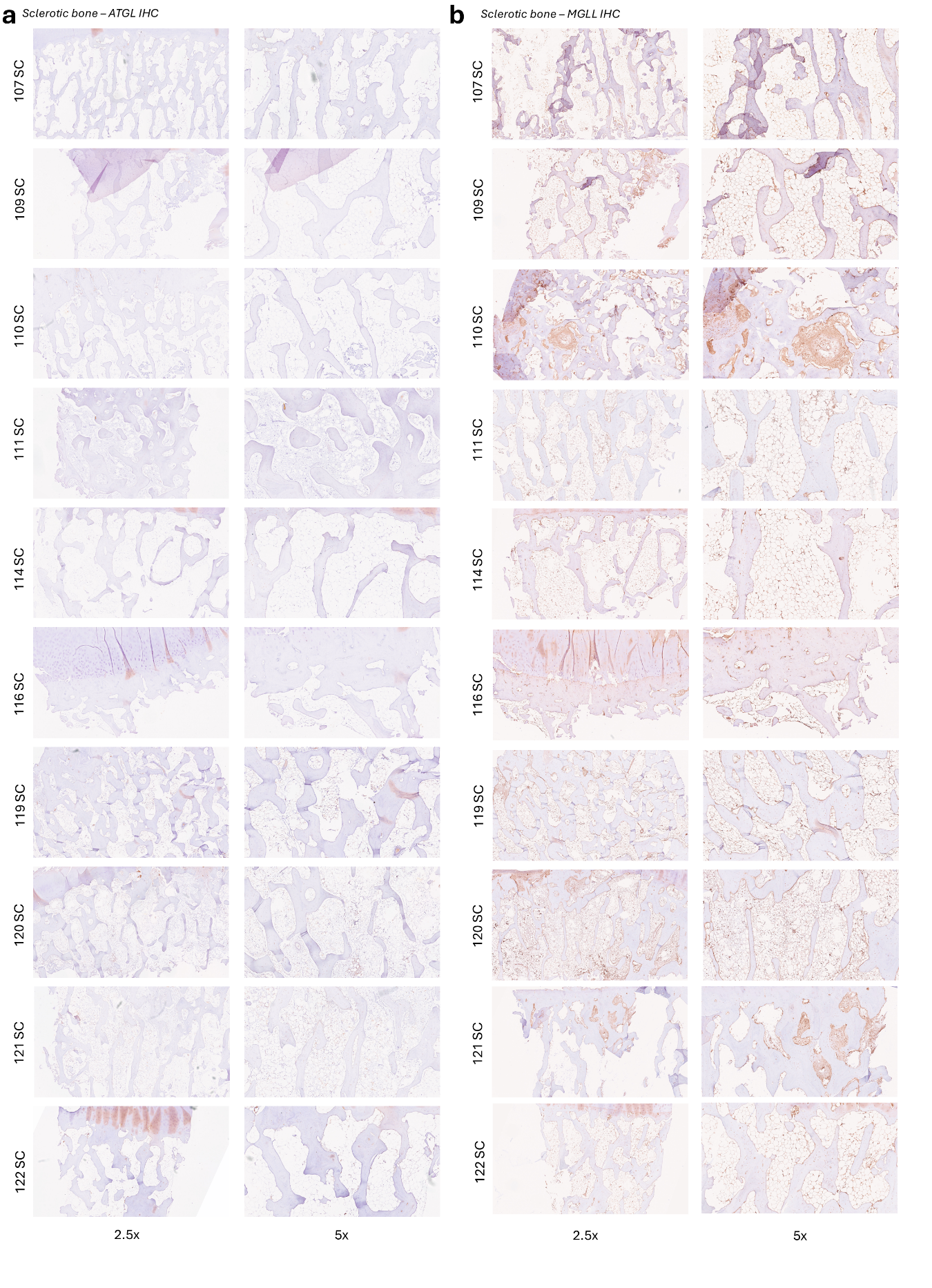

Immunohistochemical staining for (a) ATGL and (b) MGLL in sclerotic subchondral bone samples from all donors included in the analysis. Images are shown at 2.5× and 5× magnification. Brown staining indicates immunoreactivity, with hematoxylin counterstaining.

**Supplementary Figure 4. Immunohistochemical staining for HSL and phosphorylated HSL in sclerotic subchondral bone**

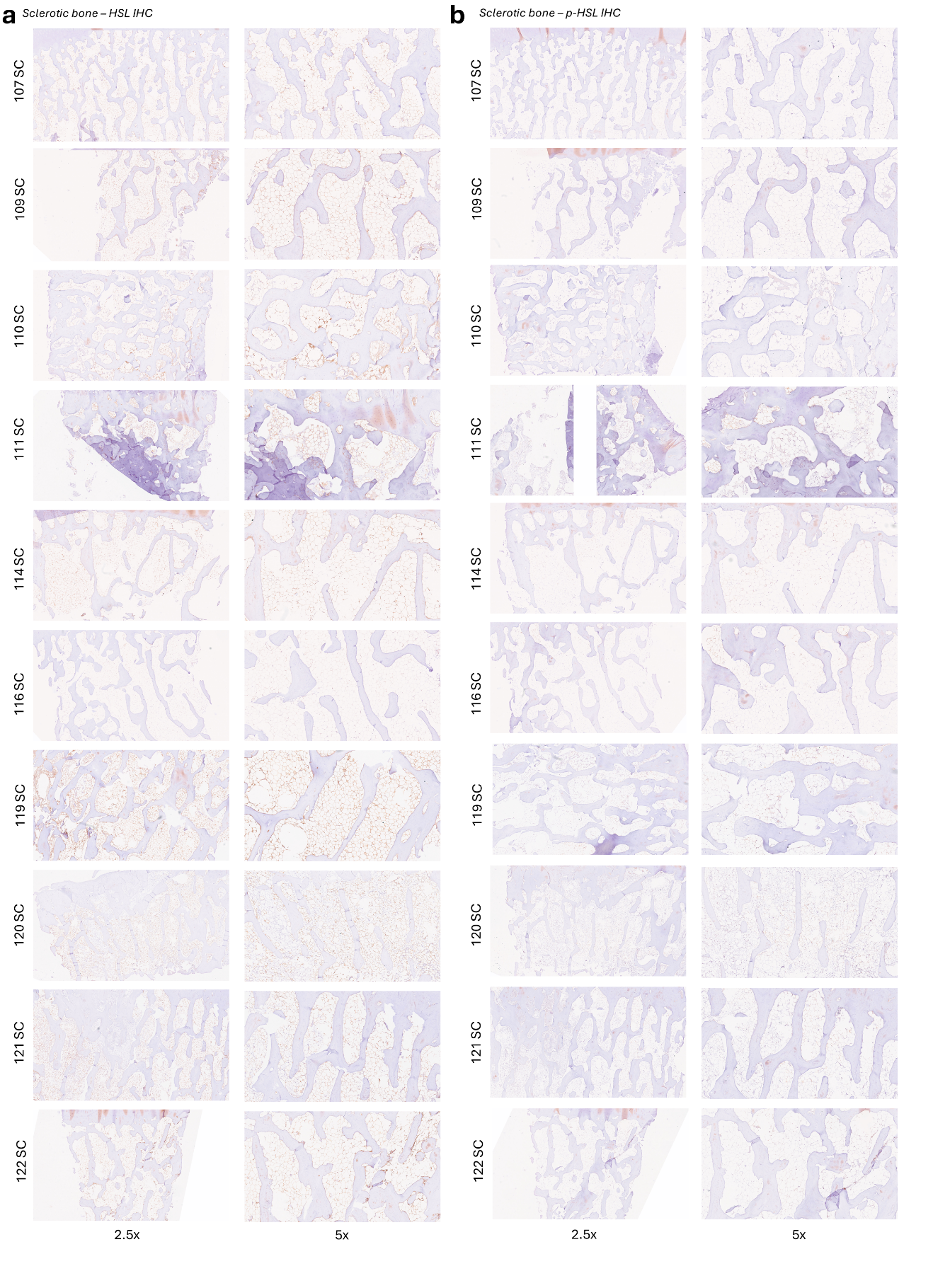

Immunohistochemical staining for (a) HSL and (b) (p-HSL) in sclerotic subchondral bone samples. Images are shown at 2.5× and 5× magnification. Brown staining indicates immunoreactivity, with hematoxylin.	counterstaining.
